# Development of an AI Algorithm for Automatic Classification of Gram Stain Images in Microbiology

**DOI:** 10.64898/2025.12.03.692028

**Authors:** Serghei Musaji, Pamela Kibsey, Andrei Musaji

**Affiliations:** New Brunswick University; Medical Microbiology Laboratory, Royal Jubilee Hospital, Island Health, British Columbia, Canada

**Keywords:** Gram stain images, microbiology, artificial intelligence, image processing, contour detection, machine learning, random forest algorithm, image classification, algorithm development

## Abstract

This paper reflects on the development and performance of an advanced artificial intelligence (AI) algorithm for the automated processing and classification of Gram stain images obtained from actual microbiology samples used in clinical microbiology. The aim of the project was to effectively categorize non-standardized Gram stain images into the six most common categories: Gram-negative rods, Gram-positive cocci in chains, Gram-positive cocci in clusters, Gram-positive rods, Gram-negative cocci, and yeasts. The development and testing relied on 1,077 Gram stain images of varying sizes, originating from different laboratories and captured using diverse microscopes at different points in time, resulting in differences in image quality, scaling, color balance, and the presence of artifacts. The dataset was split into 80% training and 20% testing subsets, with the split performed in a stratified manner so that each object group was proportionally represented in both the training and testing sets. Preprocessing involved computer vision techniques to improve contrast and color balance, detect contours and object borders, and implement filtering mechanisms to remove unwanted artifacts. Morphological analysis of shapes was then performed to extract parameters characterizing each contour. Next, human-like classification criteria—based on gradient, morphological features (e.g., shape, size) and spatial arrangement that mimic microbiologists’ visual assessment—were established, achieving around 92% accuracy in image classification without using machine learning (ML) methods. However, any further improvements turned practically impossible, prompting the use of ML methods. Building on pre-obtained features, a random forest ML algorithm was employed to further refine the criteria, with three models trained and tested successively. The first model determined the Gram stain reaction (positive or negative) of each object. The second model classified objects into one of six predefined categories. The third model aggregated individual object classifications to generate an overall classification for each slide, based on the number of objects observed in each category and their occupied area. Overall, the ML solution was significantly more accurate, reaching 99.9% accuracy in classifying the images into one of the aforementioned groups. The algorithm’s limitations include inability to classify mixed cultures, as it primarily focuses on the dominant category. In cases where positive and negative objects coexist, the algorithm tends to prioritize Gram-positive objects. Additionally, the current morphological assessment is insufficient for yeast classification. Addressing these limitations is a crucial avenue for future research to enhance the algorithm’s versatility and accuracy.

## BACKGROUND

Gram staining is a fundamental technique in microbiology (1) used to differentiate bacteria into two major groups—Gram-positive and Gram-negative—based on differences in cell wall composition. Beyond this primary distinction, microorganisms can be further classified by morphology, such as cocci, rods, or yeasts. Spatial arrangements, including cocci in chains or clusters, provide additional diagnostic clues with significant clinical relevance. The Gram staining method is routinely employed in hospitals, clinics, and diagnostic laboratories worldwide. Accurate classification of microbial species is critical for diagnosis, as it directly influences antibiotic selection and treatment strategies. It also supports research by facilitating studies of bacterial transmission, mutation, and drug resistance. Gram-stained slides additionally serve as an important tool for quality control in microbiological laboratories. They are routinely used to verify culture purity and to assess the morphology of bacterial isolates. However, traditional final evaluation relies on manual interpretation by trained microbiologists—a process that is time-consuming, subjective, and prone to variability. Differences in staining protocols, slide preparation, and individual interpretation can significantly affect both accuracy and consistency.

Despite being developed more than a century ago, the Gram stain remains a rapid, inexpensive, and indispensable technique that supports initial diagnostics and guides subsequent laboratory testing, including culture-based assays, biochemical profiling, and molecular methods such as PCR and sequencing (2). Nevertheless, the simplicity and low cost of Gram staining have resulted in relatively limited attention from artificial intelligence and machine learning (AI/ML) researchers, leaving a clear gap in the field. Yet these same characteristics make the method particularly valuable in under-resourced settings, where AI-assisted analysis could enable minimally trained personnel to make reliable treatment decisions using basic microscopy. Developing effective AI/ML systems for Gram stain interpretation, however, remains challenging even when high-quality imaging equipment is available. Most existing approaches depend on standardized staining and imaging conditions—requiring specific microscope– camera combinations, fixed magnifications, and Whole Slide Imaging—to ensure consistent data quality.

In contrast, our work focuses on developing a more flexible algorithm with low sensitivity to variations in image quality. Such variations commonly arise from inconsistent staining, differing equipment, and variable microscope calibration or maintenance (3). This project, therefore, addresses the need for cost-effective, efficient, accessible, and objective analytical tools in microbiology. Here, we present the initial steps in the design and implementation of an algorithm capable of accurately processing and classifying diverse, real-world laboratory images. Ultimately, this approach could enable diagnostic applications in resource-limited environments—for example, allowing images captured through a smartphone camera positioned over a conventional analog microscope to be analyzed in mobile or field hospitals using online or off-line AI/ML algorithms. This approach may allow image acquisition and access to analysis around the World, including resource-limited settings (4)

## METHODOLOGY

### Image set

To address our objective, we collected a sample of 1,077 completely nontraceable to the primary clinical source and deidentified monoorganismal blood culture Gram stain images sourced from various Island Health, BC, Canada microbiology laboratories. The images originated from routine operations and, thus, exhibited variability in size, microscopes used, focus quality, color balance, and the presence of artifacts. The images were accompanied by the information on their classification by the lab technician. All slides were further verified and validated by at least two professional microbiologists, who had to agree on the Gram stain classifications. Our aim was to develop an algorithm to replicate microbiology technologist-physician decisions. Type of images used for this study are shown in Table 1 with examples presented in Table 2.

**Table 1.** Gram stain image groupings used in this study.

| Slide type | Slide count |
| --- | --- |
| Gram Positive Cocci in Chains | 247 |
| Gram Positive Cocci in Clumps | 306 |
| Gram Negative Rods | 244 |
| Gram Positive Rods | 192 |
| Gram Negative Cocci | 25 |
| Yeasts | 63 |
| **Total** | **1077** |

**Table 2.**
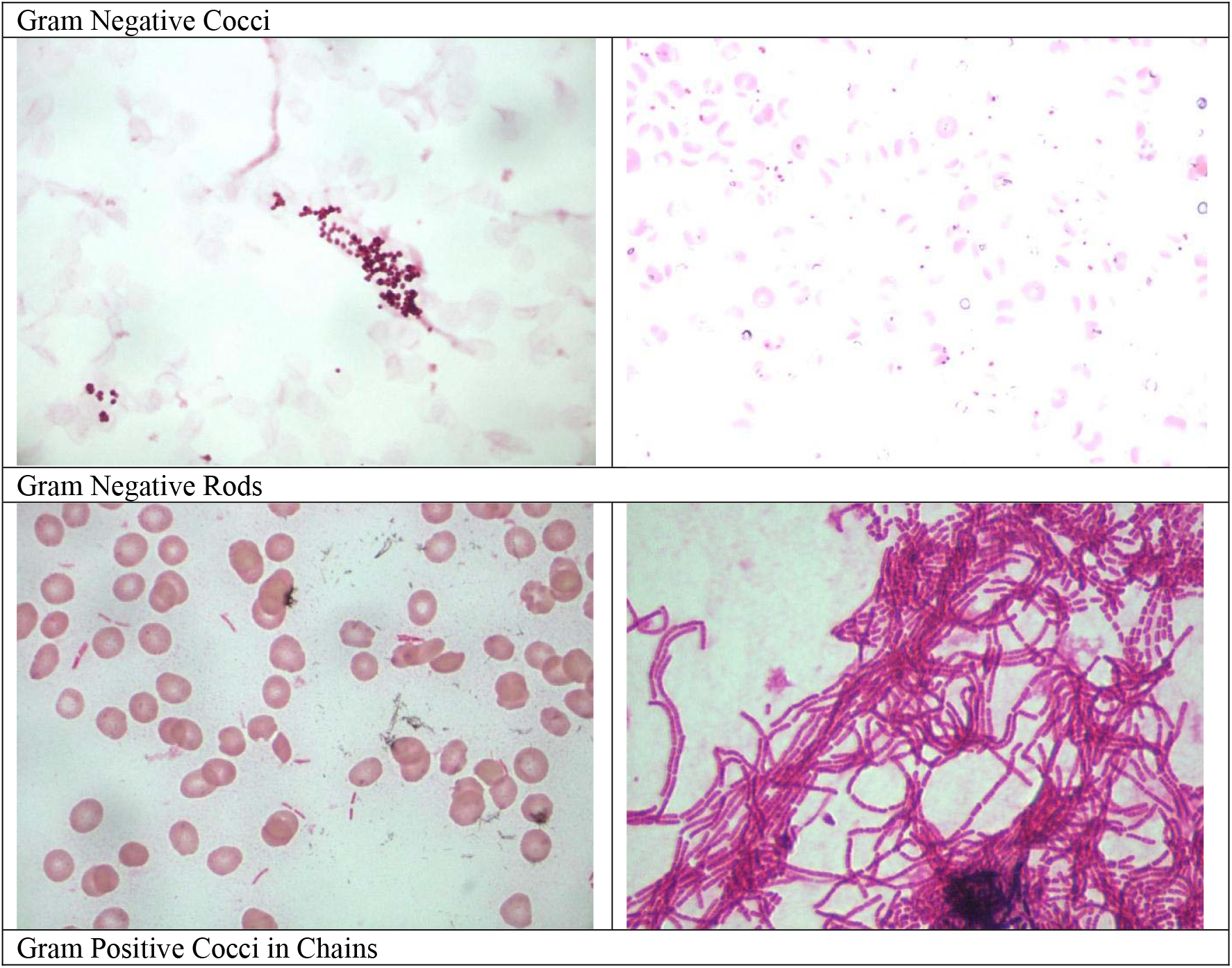
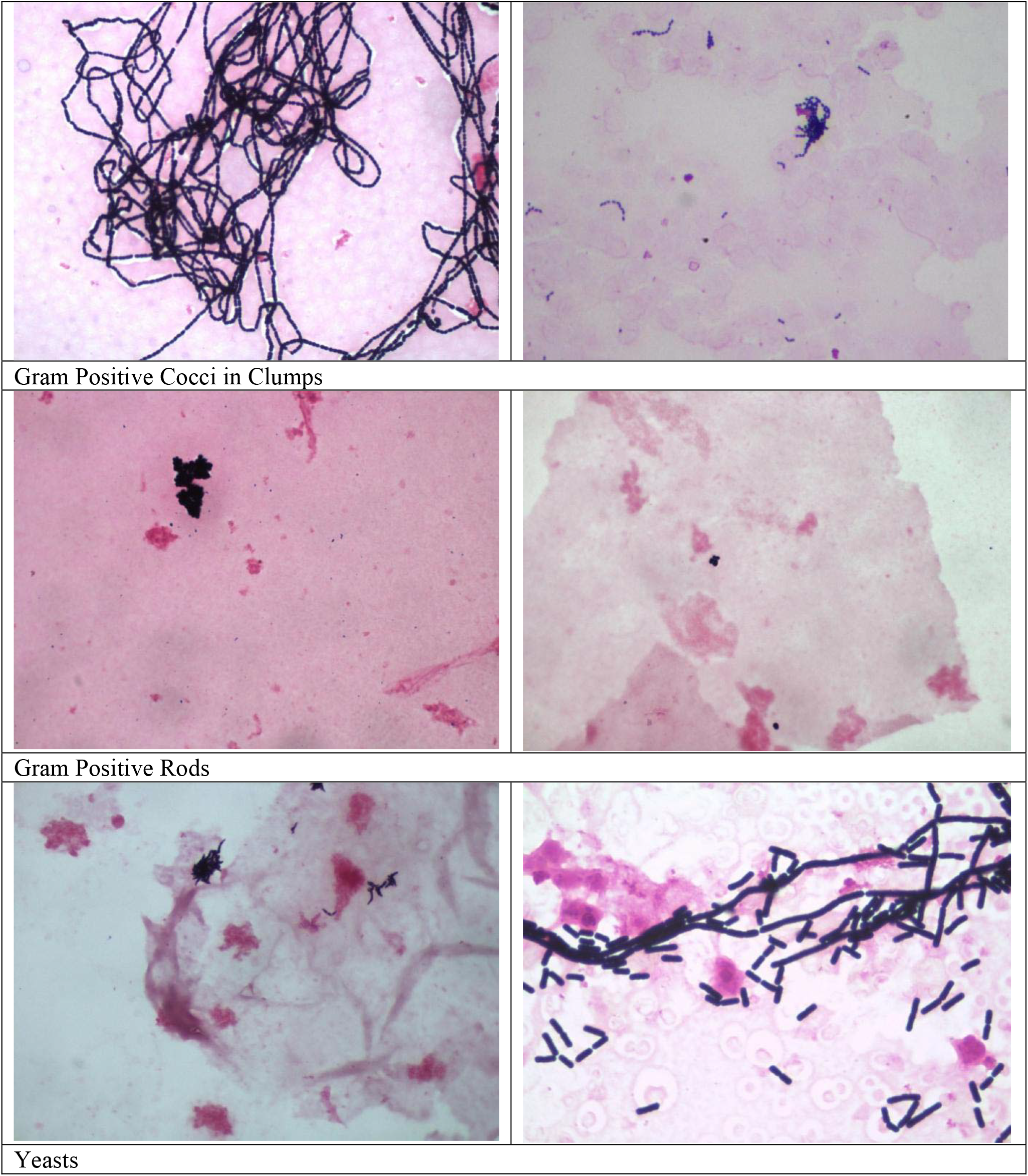
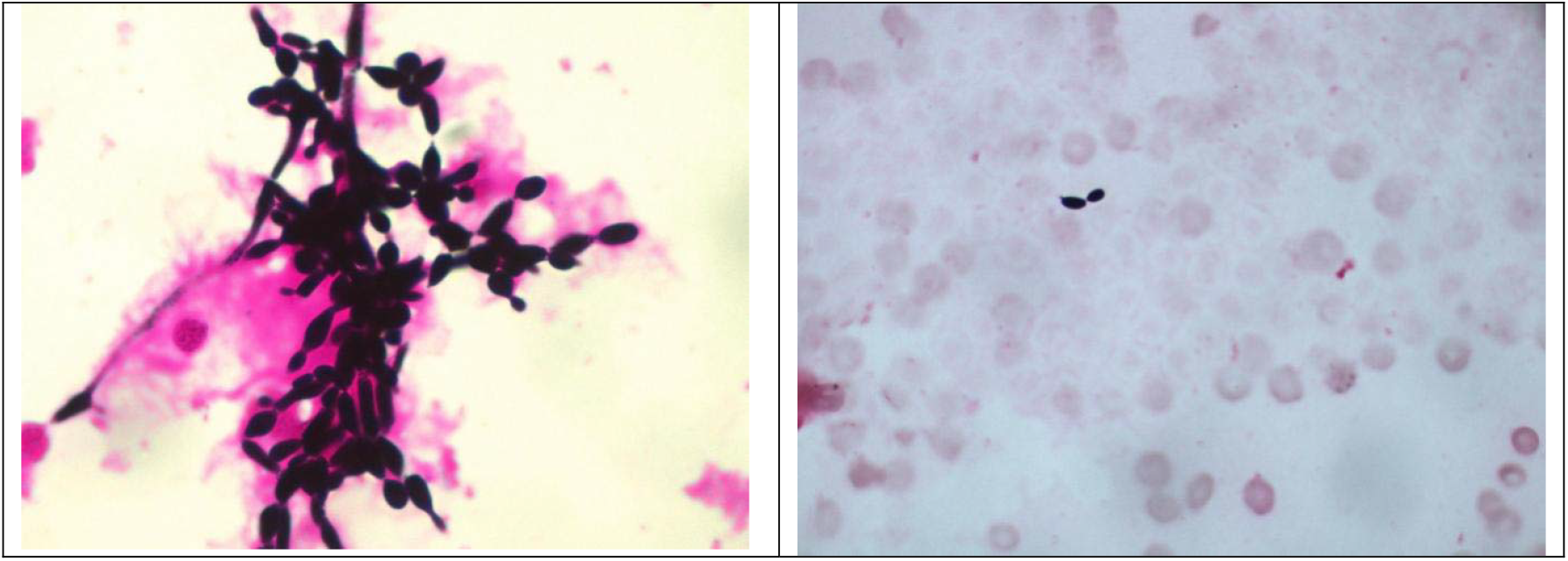
Examples of studied images.

### Algorithm development

To prepare the images, computer vision techniques were applied to enhance sharpness, contrast, and brightness, improving contour detection in subsequent stages. Contour detection algorithms were then used to identify object borders, providing the foundation for isolating individual bacterial or fungal structures. A filtering step removed unwanted artifacts and extraneous objects—ranging from erythrocytes to marks caused by dirty lenses—ensuring cleaner datasets and more reliable downstream analyses.

Morphological analysis was subsequently performed to extract parameters characterizing each contour, including length, width, area, curvilinearity, indentations, and dominant color, among other relevant parameters. These quantitative features formed the basis for classification. Initial attempts employed manually programmed, rule-based criteria designed to replicate human analytical reasoning. Using this approach, Gram-positive versus Gram-negative differentiation achieved over 99% accuracy, even without artificial intelligence. Further manual programming enabled classification of individual objects and slides according to bacterial morphology (e.g., length-to-width ratio, curvilinearity, indentation), color characteristics (e.g., intensity and gradient), and spatial organization (e.g., chains versus clusters), reaching approximately 92% accuracy. To visualize this process, we further present some examples of processed slides with some annotations and classification decisions based on the manually set criteria (See Images 1 & 2).

**Image 1.**
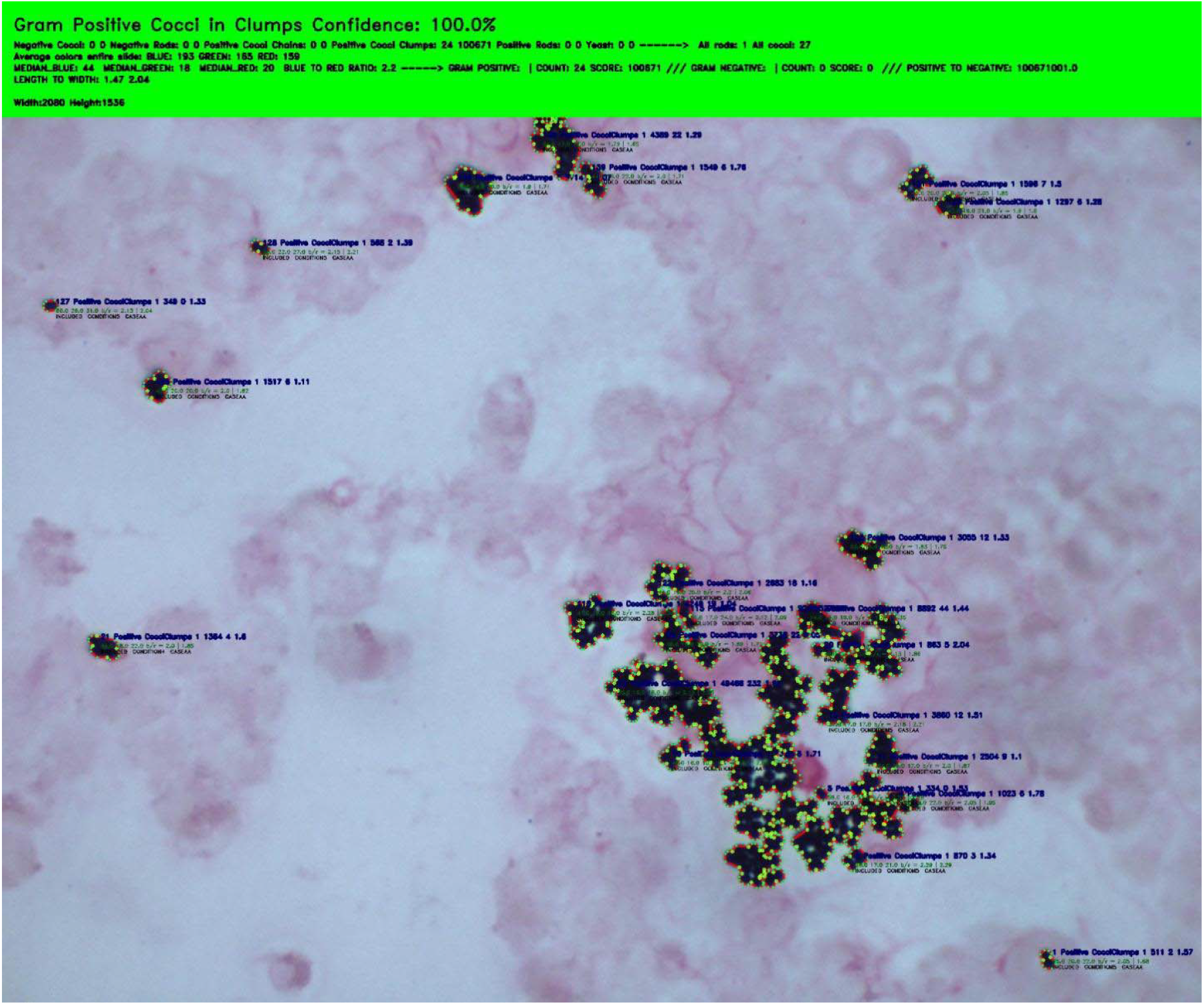
Classification based on manually set criteria – Gram Positive Cocci in Clumps.

**Image 2.**
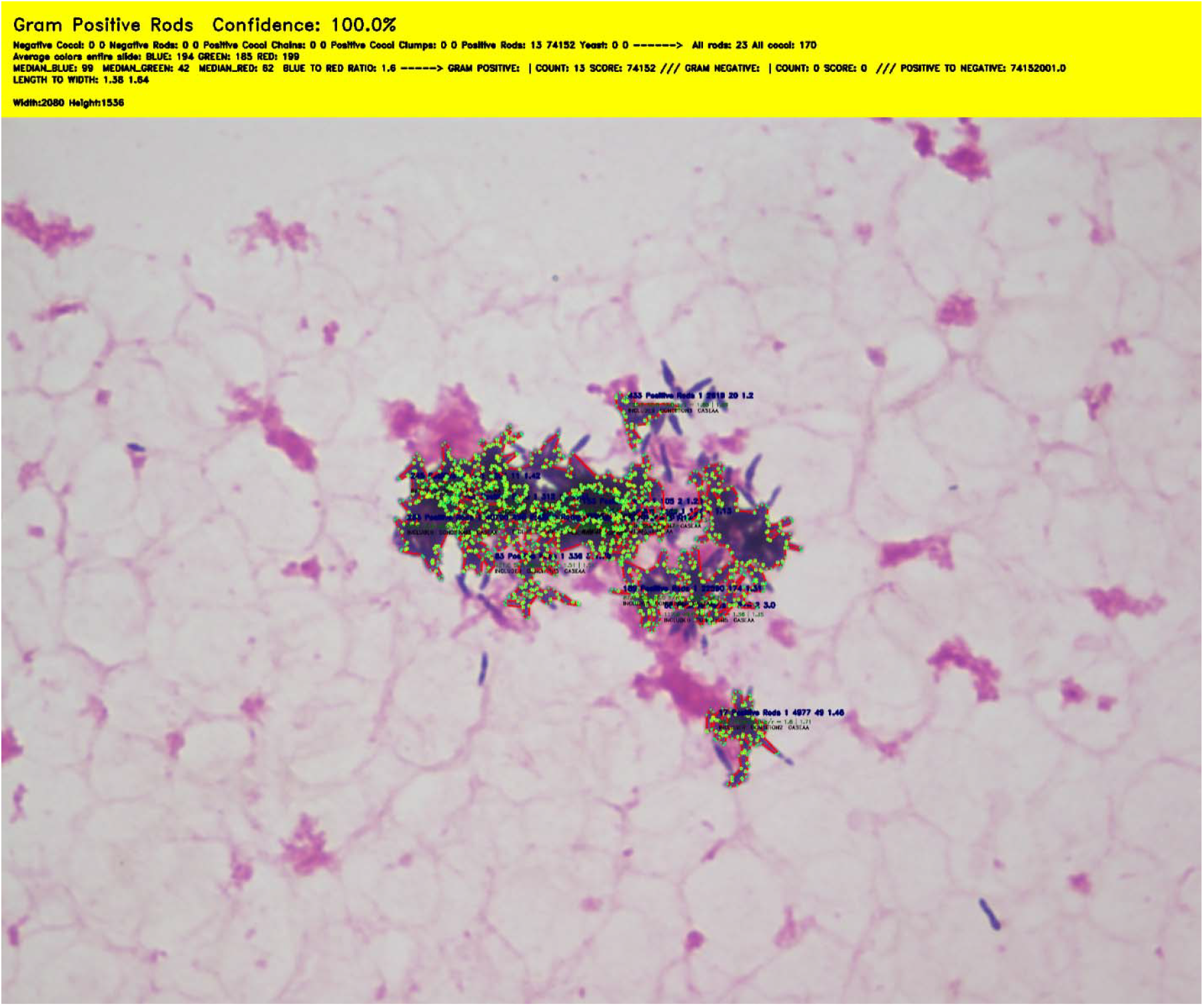
Classification based on manually set criteria – Gram Positive Rods.

Unfortunately, the increasing complexity and variability of bacterial patterns limited the potential for further improvement with purely manual methods.

To address these limitations, machine learning (ML) optimization was introduced, specifically using a random forest algorithm. Random forests were selected due to their suitability for small datasets, computational efficiency, and high performance on structured, feature-based data. Unlike convolutional neural networks (CNNs), which require large, labeled image datasets and significant computational resources, random forests can achieve excellent results with modest hardware and limited sample sizes.

Rather than manually annotating each bacterial object, we extracted a comprehensive set of quantitative image descriptors—including color histograms, intensity gradients, texture metrics, and geometric or morphological parameters—and trained the model to classify based on these features. Random forests effectively capture nonlinear relationships and interactions among multiple quantitative variables while remaining resistant to overfitting. Their ensemble nature, combining multiple decision trees, ensures robustness against noise and variability, which are common in Gram-stain images due to differences in staining quality, lighting, and microscope calibration. CNNs, in contrast, would have required strict image standardization, larger datasets, and greater computational power, making them impractical in this study. Thus, the random forest approach provided a practical, scalable, and interpretable solution.

For model training, the dataset was divided using an 80–20 stratified partitioning strategy, allocating 80% of slides to the training set and 20% to the testing set, while ensuring proportional representation of all object groups in both subsets. Three separate models were developed to enhance accuracy and versatility. The first model distinguished Gram-positive from Gram-negative images. The second model identified individual objects within the images. Finally, a third model classified each slide into one of six predefined groups, based on the number of detected objects in each class, their relative total area, and the spatial distribution of relevant objects within the image.

## RESULTS

The ML-based algorithm achieved an impressive accuracy rate of 99.7%, representing a substantial improvement over the initial manual classification criteria, which achieved approximately 92%. Each of the three models was evaluated individually to assess its specific contribution to the overall algorithm’s performance.

Model 1, which focused on distinguishing Gram-positive from Gram-negative images, showed only marginal improvement with ML optimization. Model 2, responsible for category-level classification, demonstrated the most pronounced increase in accuracy. Model 3, designed to analyze object arrangement and spatial organization, further highlighted the adaptability and efficiency of ML-based decision-making processes.

The final classification results are summarized below. Only a single slide was misclassified (see Image 3). Expert evaluation identified this slide as containing Gram-positive cocci in clumps, whereas the algorithm classified it as Gram-positive cocci in chains. This discrepancy likely arises from the behavior of the computer vision and contour detection algorithms, which outlined features resembling linearly arranged cocci—interpreted as chains. Because cocci in chains often overlap, producing spatial patterns visually similar to clumps, both manual and ML-based classification systems may interpret such configurations as stronger indicators of chain-like arrangements rather than true clumping.

**Image 3.**
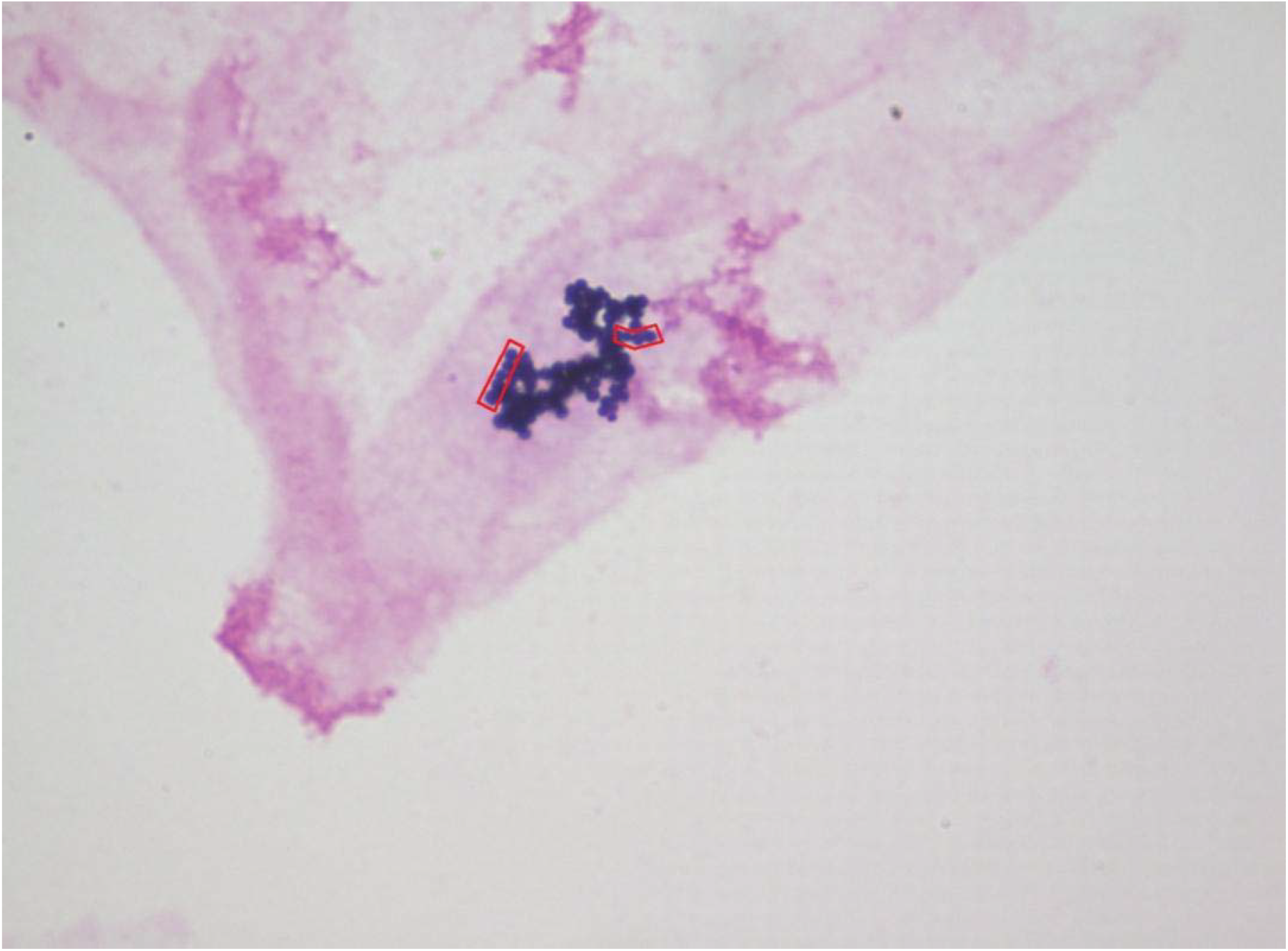
Misclassified slide. Classified by our AI as Gram Positive Cocci in Chains, whereas classified by the experts as Gram Positive Cocci in Clumps.

In fact, it is not a surprise that the algorithm ended-up classifying this slide as containing Gram Positive Cocci in chains since in a few other instances, similar-shaped objects were classified by the experts as being Gram Positive Cocci in chains (see Image 2).

Image 4 clearly illustrates the presence of Gram-positive cocci in chains, primarily because the slide contains multiple, well-defined chain formations. In contrast, Image 1 includes only a single object, making classification inherently more challenging.

**Image 4.**
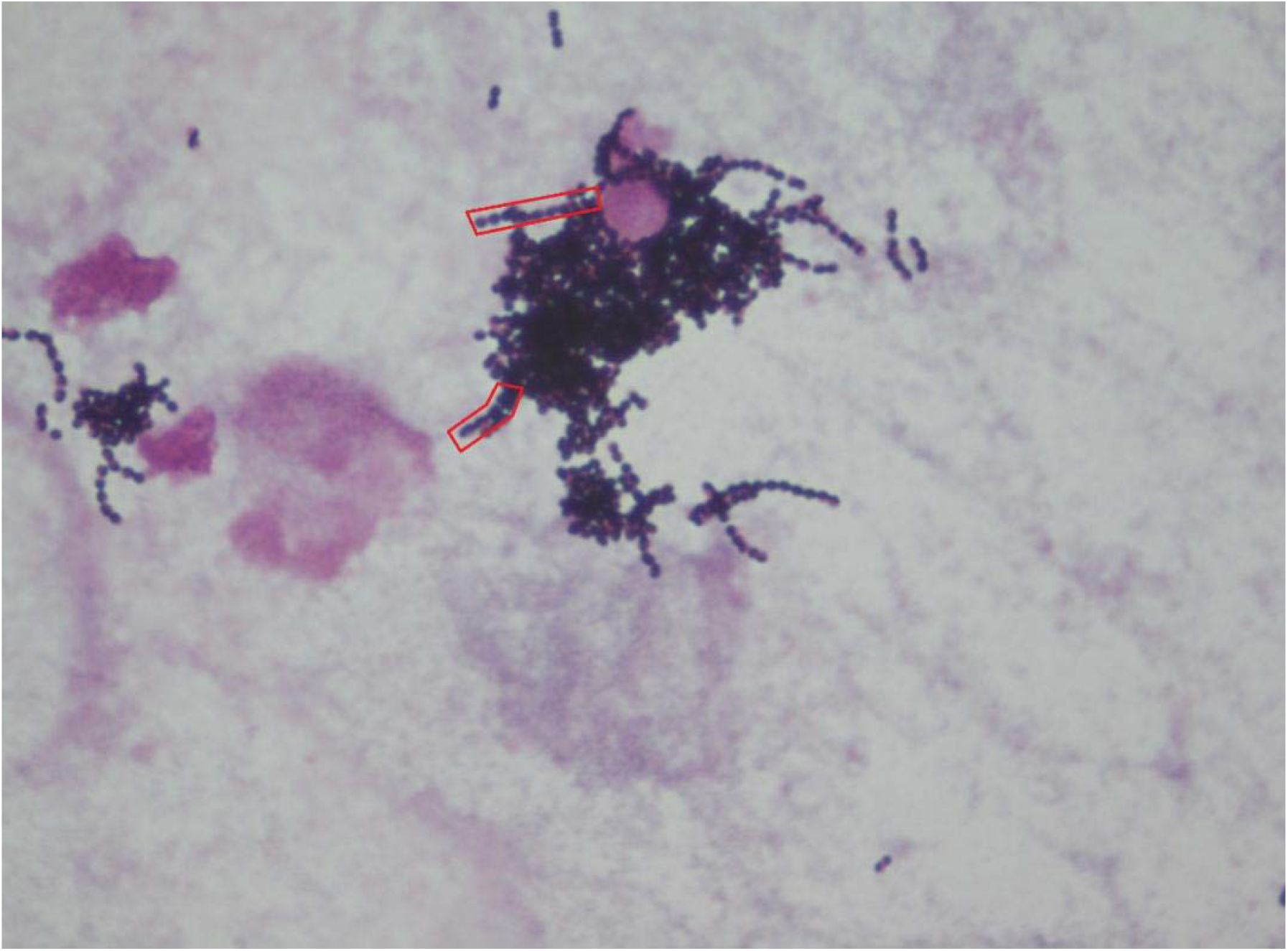
Classified by AI and the experts as Gram Positive Cocci in Chains.

Data comparing expert and AI/ML-based image categorizations are summarized in Tables 3 and 4. As shown, the AI algorithm performed consistently well across all categories, achieving Gram-stain classification accuracy rates above 95%—and reaching 100% in most categories— except for Gram-positive cocci in clumps, which exhibited a slightly lower accuracy of 99.7%.

**Table 3.**
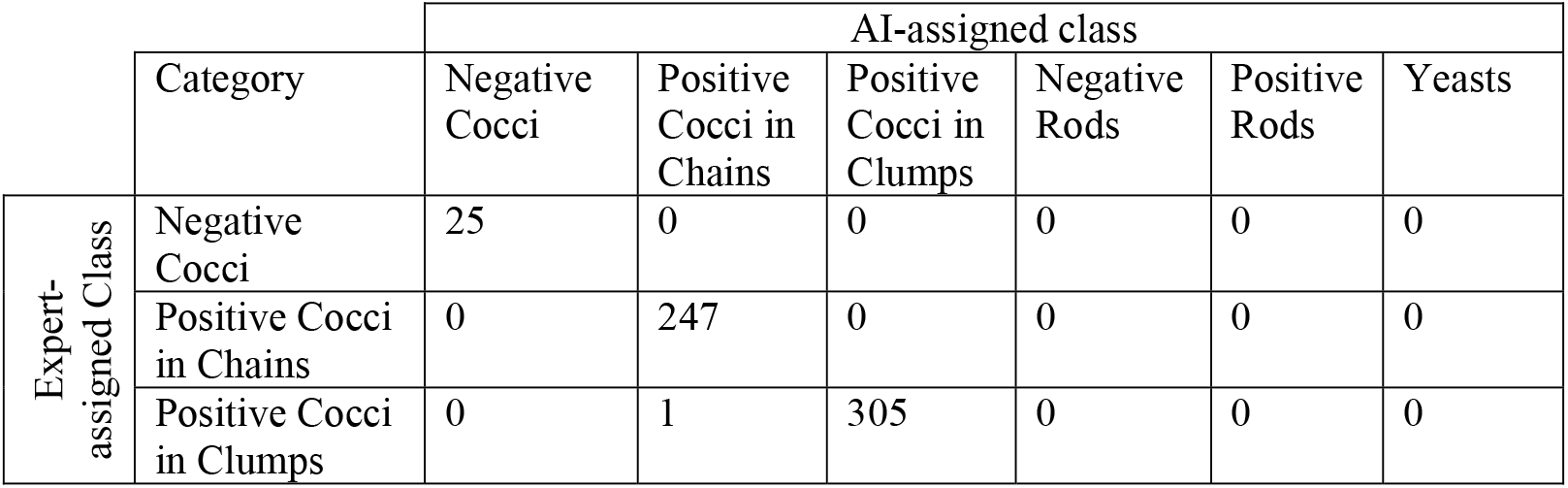
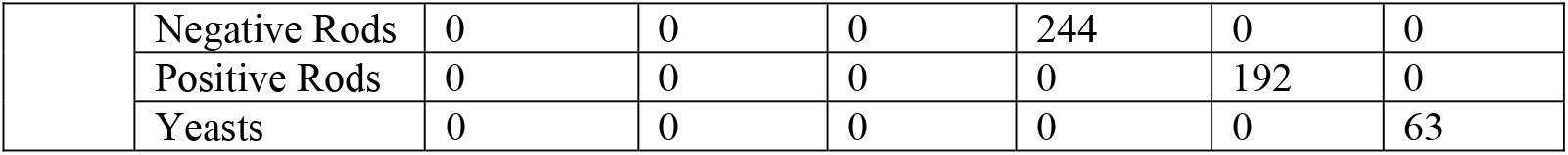
Classification of images into different classes (slide count) by AI vs expert-defined (gold standard comparator)

**Table 4.**
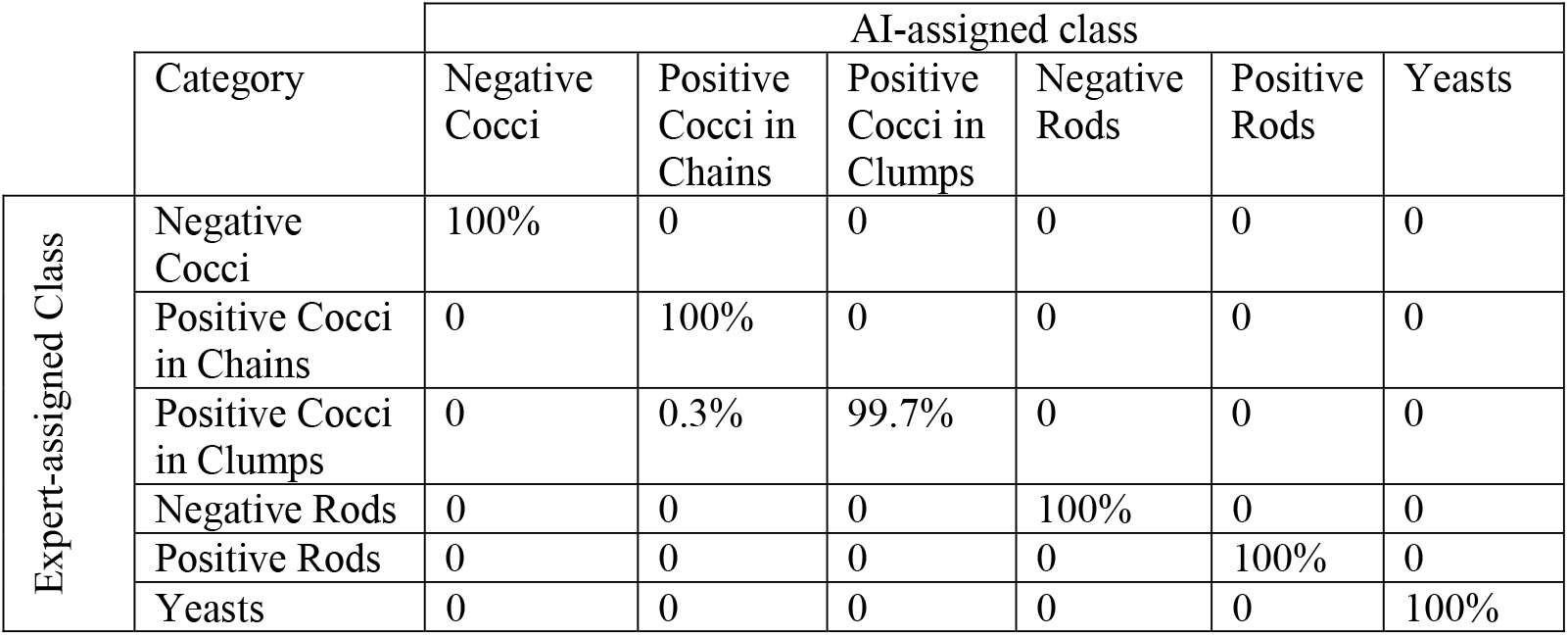
Accuracy of Classification of images into different classes (percentages) by AI vs expert-defined (gold standard comparator)

## DISCUSSION AND IMPLICATIONS

The development of an AI-driven algorithm for the automatic classification of Gram-stained images represents a significant advancement in microbiological diagnostics and research. Built on a foundation of robust image-processing methods and optimized using machine learning (ML)—specifically, the random forest algorithm—the system demonstrates exceptional performance in real-world laboratory conditions. While the manually programmed approach achieved accuracy around 92%, the integration of ML elevated overall accuracy to 99.9%, underscoring the effectiveness of combining computer vision with data-driven learning techniques.

Beyond its high accuracy, the AI algorithm offers clear benefits in cost-effectiveness, efficiency, and accessibility. In resource-limited settings—where advanced imaging systems and trained microbiologists may be scarce—the algorithm could provide a low-cost alternative for reliable Gram-stain interpretation. By automating image classification, it minimizes dependence on expert judgment and expensive equipment, thereby democratizing access to accurate diagnostic tools and improving healthcare outcomes in underserved regions.

Automation also enhances laboratory efficiency. Traditional manual interpretation of slides is time-consuming and highly dependent on human expertise. The AI algorithm standardizes and accelerates the classification process, enabling microbiologists to focus on higher-level diagnostic reasoning and research. This improvement translates into faster turnaround times, reduced operational costs, and increased productivity in both clinical and research environments.

Furthermore, the algorithm contributes to standardization and consistency in microbiological practice. Human interpretation is inherently variable, particularly across institutions with differing staining protocols and microscopy standards. The ML-based approach minimizes such inconsistencies, producing objective, reproducible classifications that can be compared across laboratories. Standardized digital workflows also support the establishment of reference datasets and benchmarks, strengthening quality assurance in microbiological diagnostics.

The algorithm’s digital and automated nature also facilitates remote and collaborative applications. Digitized Gram-stain images can be analyzed and shared across laboratories, with the algorithm providing consistent, objective classifications regardless of location. This capability supports telemicrobiology, remote consultation, and cross-institutional research collaborations, ultimately contributing to global infectious disease monitoring and diagnostic capacity building.

From a commercial standpoint, the algorithm holds strong potential for integration into existing digital microscopy and diagnostic imaging platforms. It could be implemented as a standalone software solution, integrated into microscope camera systems, or deployed via cloud-based services to enable automated Gram-stain analysis. Such implementations could appeal to diagnostic equipment manufacturers, telehealth providers, and laboratory automation companies seeking cost-effective AI-enhanced tools.

As the algorithm continues to evolve through larger, more diverse datasets and iterative refinement, its reliability and commercial readiness are expected to grow. Ultimately, the system’s combination of high accuracy, adaptability, and scalability positions it as a valuable tool for advancing microbiology and laboratory diagnostics worldwide.

### LIMITATIONS AND FUTURE RESEARCH

Despite the algorithm’s success, several limitations must be acknowledged. One of the main challenges lies in its inability to accurately classify mixed cultures and its tendency to prioritize dominant categories when both Gram-positive and Gram-negative organisms are present. Although mixed blood cultures are relatively uncommon, addressing this limitation is critical for improving diagnostic reliability and expanding the algorithm’s clinical utility.

The dataset size also presents a constraint. The number of training samples was limited, with relatively few observations for certain classes—particularly yeast and Gram-negative cocci—which may have affected the model’s ability to generalize across all microbial categories. Moreover, while the images were obtained from different laboratory microscopes, the overall pool of equipment was relatively narrow. Although this helped maintain procedural consistency, it may have inadvertently constrained the algorithm’s adaptability to wider variations in image acquisition settings.

Image quality variability further influenced performance. Some slides exhibited poor staining or imaging artifacts, which provided valuable insights into algorithmic robustness; however, the technicians generally followed similar staining protocols, resulting in a relatively uniform dataset. This homogeneity, while beneficial for model training, may limit generalizability to data acquired under significantly different conditions—such as alternative staining reagents, microscope optics, or lighting setups. It remains to be seen whether the algorithm can adapt effectively to greater variance in scaling, and color balance, particularly in low-resource settings using analog microscopes or mobile phone–based imaging systems.

Future research should therefore focus on enhancing the algorithm’s capacity to handle mixed cultures and mitigate prioritization bias in heterogeneous samples. Expanding the dataset to include a broader range of microorganisms, imaging equipment, and staining conditions will be essential for improving model robustness and generalizability. Additionally, exploring other ML architectures—including ensemble methods and deep learning approaches—may further enhance performance and adaptability.

Finally, it would be valuable to benchmark specialized Gram-stain classification algorithms against broader, general-purpose AI systems such as ChatGPT or multimodal vision-language models to evaluate their relative interpretive and decision-making capabilities. It would also be informative to test the algorithm under deliberately degraded imaging conditions to assess its resilience and practical usability in resource-limited environments.

